# High nutrient loads hinder successful restoration of natural habitats in freshwater wetlands

**DOI:** 10.1101/2022.03.10.483603

**Authors:** Jesper Erenskjold Moeslund, Dagmar Kappel Andersen, Ane Kirstine Brunbjerg, Camilla Fløjgaard, Bettina Nygaard, Rasmus Ejrnæs

## Abstract

Restoration of natural processes in ecosystems is key to halt the biodiversity crisis. Here, we evaluate 20 different stream-valley wetland restoration projects – mainly rewetting – in a large region in Denmark in terms of successful restoration of natural wetland habitats. We used quadratic discriminant analysis and generalized linear models to compare the projects’ 80 vegetation plots with >50.000 natural wetland-habitat reference plots and modelled the influence of time, grazing, rewetting and nutrient availability on the study plots’ probabilities of belonging to such natural habitats and their richness of high-quality habitat indicator species. In our study, the probability of a restored wetland being a natural wetland habitat – almost always an alkaline fen – was generally below 10 %. Also, we only found half as many indicator species in restored wetlands than in reference wetlands and we demonstrated that the number of characteristic alkaline fen species did not deviate from what could be expected under the prevailing nutrient conditions. We found a negative effect of nutrient availability on the number of high-quality habitat indicator species and the lowest probability of plots being natural wetlands in the most nutrient rich plots. The effect of grazing was only positive in the first years after restoration and only in the most nutrient rich plots, while the effect of rewetting sites to their historical hydrological conditions was generally negative. Our findings reveal that unnaturally high nutrient availability is probably the core limiting factor for successful restoration of natural wetlands and their associated plant diversity.

**Implications for practice:**

- To successfully restore natural and characteristic freshwater wetland habitats focus on recreating natural processes and conditions is needed
- Restoring natural hydrology and grazing is not enough, the soil and water must be naturally nutrient poor for successful restoration of these habitats
- Restoration of stream-valley wetlands such as alkaline springs and fens is more likely to be successful in spring-dominated landscapes where clean groundwater diffusely exfiltrates the soil

## Introduction

Freshwater wetlands are vital ecosystems for the conservation of biodiversity. They are sensitive to even small land-use changes and they hold a high number of red-listed species (Moeslund et al., 2019; Tickner et al., 2020). In modern times, 54–57 % of wetlands globally have been lost and since 1700 AD this number is probably up to 87 % (Davidson, 2014). Many freshwater wetlands have been lost or degraded as a consequence of altered hydrology due to drainage, drinking water extraction, dam-building and other human-made changes to the natural water cycle (Brinson & Malvárez, 2002; Rolls et al., 2018). Also, in the last century the natural disturbances by large herbivores – e.g., maintaining open vegetation and leaving dung (and in some cases carcasses) – have dramatically decreased in these important ecosystems, further contributing to biodiversity losses (Svenning et al., 2016; Biró et al., 2019). Here, we evaluate 20 different cross-regional restoration projects (mainly hydrology) in Denmark to point out the most important factors for ensuring cost-effective restoration of natural wetland habitats.

In Northern Europe, stream-valley (i.e., riparian) wetlands typically count habitats such as alkaline fens, Molinia-meadows, and spring-communities. North European alkaline fens are characterised by the seepage of base-rich groundwater, with several *Carex* species and e.g., *Schoenus nigricans* L., *Juncus subnodulosus* Schrank, *Liparis loeselii* (L.) Rich. and *Pinguicula vulgaris* L. The often surface-water-affected Molinia-meadows typically hold *Molinia caerulea* (L.) Moench, and several *Juncus* species e.g., *Juncus conglomeratus* L. (European Commision, 2013). Finally, spring communities can vary in species composition depending on the soil characteristics and openness of the vegetation. Natural disturbances such as groundwater seepage, flooding and grazing combined with a naturally nutrient poor and base-rich environment has historically caused these wetland communities to be rather open swards and rich in plant and bryophyte species (Bergamini et al., 2001; Moran et al., 2008; Biró et al., 2019). Consequently, they have been listed by the European Union as important habitats for the European community to conserve and protect within their natural ranges (Council of the European Communities, 1992).

Globally, efforts to restore freshwater wetlands to their former extents and conditions are widespread (Perry, 2004; Mälson et al., 2008; Menichino et al., 2016). However, most often restoration has not aimed solely to restore natural habitats and their associated biodiversity, but also or even primarily targeted binding CO2, downstream flood prevention or providing denitrification and phosphor removal services to lower the impact of nutrients in the recipient seascapes (Zedler, 2000; Hoffmann & Baattrup-Pedersen, 2007; Moreno-Mateos & Comin, 2010; Audet et al., 2020; Hoffmann et al., 2020). The methods used for restoration of freshwater wetlands varies significantly between projects and can include for instance the restoration of hydrology, landscape terrain, grazing and nutrient balances as well as reintroduction of typical species (Pfadenhauer & Grootjans, 1999; Matthews et al., 2009a; Richardson et al., 2016). However, while there are numerous studies of wetland restoration methods (see references above and references therein), only few freshwater restoration projects are evaluated post-restoration (Taddeo & Dronova, 2018; Baumane et al., 2021) and even fewer are systematically monitored and hence cost-effective restoration of freshwater wetlands is still poorly supported by research and empirical evidence.

In this study, we compared 80 different vegetation plots from 20 different stream-valley freshwater-wetland restoration projects in Denmark with > 70,000 vegetation plots in near-natural as well as human-influenced wetlands (17 freshwater wetland habitat types) to gain insight into what conditions are most likely to yield successful restoration of these wet habitats. More specifically, we addressed the following questions: (1) How close are the restored wetlands to comparable natural habitats with regard to plant diversity and composition? (2) What is the role of previous land-use, grazing, time since restoration and nutrient availability for restoration success?

## Methods

### Study area

In Denmark, there are several settings (i.e., combinations of former land-use and methodology) influencing the outcome of wetland restoration projects. Most projects are conducted on former extensive farmland or improved meadows in drained and ditched stream valleys or drained lakes and some on former fish-farms. These restoration projects differ in terrain, original hydrology, soil type and -chemistry, restoration methods etc. In this study, we focused on a broad range of restoration projects grouped by their former land-use: (1) groundwater-fed fish farms in groundwater-rich settings and (2) drained extensive farmland areas in low-lying riparian settings and often historically flooded by the nearby stream, typically former intensive or extensive agricultural land. For projects in the first group, we expected that the water quality is high, since this mainly consists of pure groundwater. Hence, we hypothesized that restoration on former spring-fed fish-farms could result in a relatively high habitat quality. For projects in the second group, the water for rewetting typically comes from flooding by a stream or drainpipes and consequently we expected this water to be more nutrient rich possibly affecting the restoration of habitat quality negatively. We included 20 different restored wetlands in Jutland, Denmark, equally covering former spring-fed fish-farms and former extensive farmland areas mainly influenced by surface water or water from drainpipes (Fig. 1). The 20 sites were selected to cover a large region (Eastern Central Jutland, c. 5500 km^2^), but still located in a way that allowed us to visit 2–3 sites a day. The sites were selected to cover a broad range of different restoration settings as indicated above and included for example projects that partly or mainly aimed to improve stream water quality and projects that were more aimed towards restoring stream or wetland biodiversity. We required all sites to have at least a brief description of the restoration process, e.g., former land-use and year of restoration completion.

**Figure 1.**
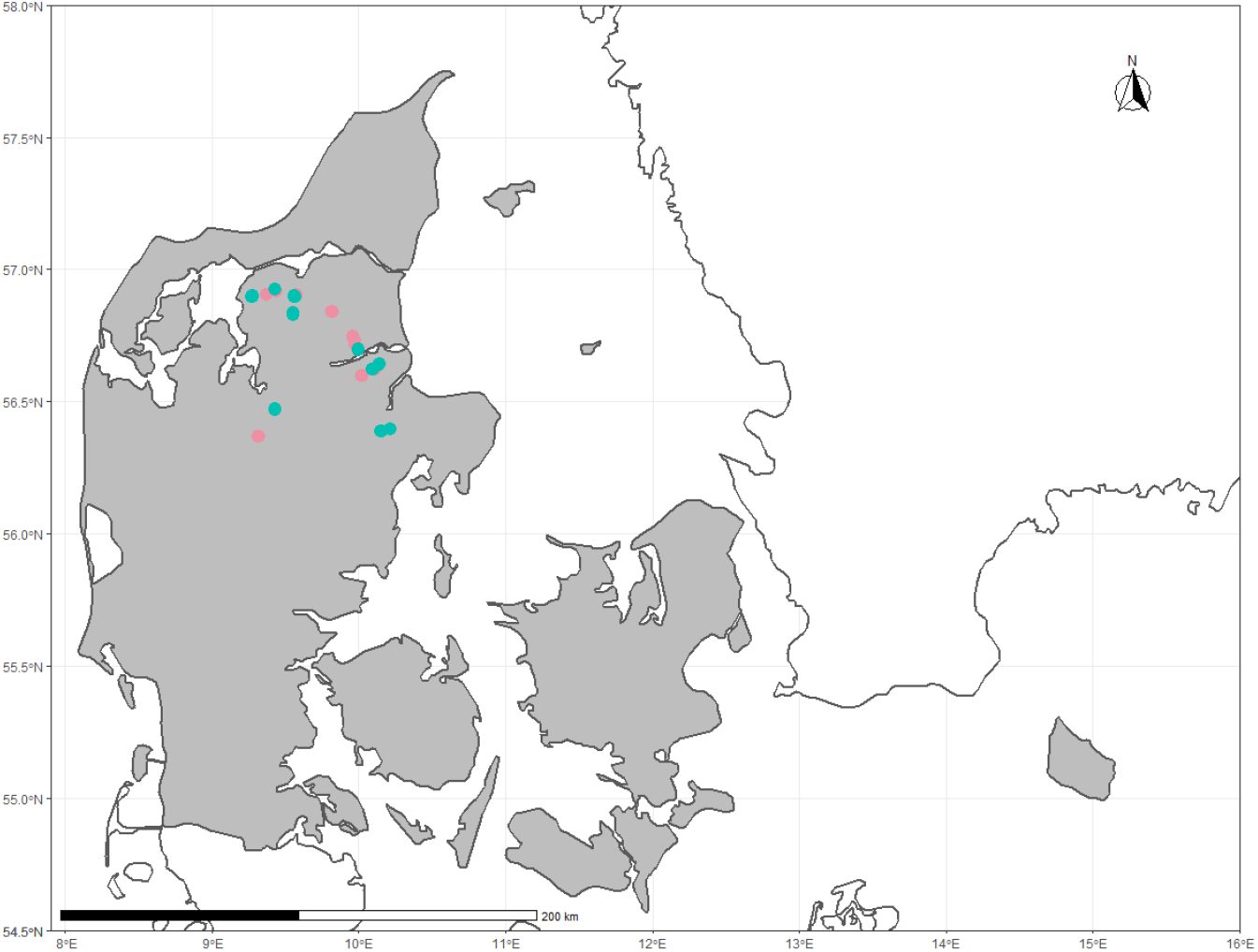
The location of our study sites in Jutland, Denmark. Turquoise plots were formerly extensive farmlands of different kind where hydrology has now been restored by reducing drainage. Light red plots were formerly spring-fed fish farms.

### Field data

In a first step, we mapped the wetness of each of the 20 restoration sites by visual inspection in the categories wet, wet-moist, moist, moist-dry and dry. The wet, moist and dry categories corresponded to the legend of a historical map that we used for modelling (see “Explanatory variables”), i.e., wet corresponded to bog or swamp, moist to meadow and dry to a dry grassland. The intermediate categories (wet-moist and moist-dry) were used in areas with mosaics of these categories. Then, in each site, we laid out four 5-m radius circular vegetation plots (i.e., 80 in total) as described in the following. If plots already existed from previous projects these were repeated (18 plots) to contribute to time series data for other studies. If not, we selected plots randomly from a 10×10 m grid avoiding plots with >50 % lake or pond coverage, >35 % woody canopy cover and plots with dry patches. We operated with these constraints as we were only interested in open terrestrial wetland habitats. Additionally, we skipped a random plot if it was <30 m away from other plots. Within each plot we recorded all vascular plant and bryophyte species. Finally, we noted whether the plot was affected by grazing by horses or cattle (i.e., bites on the vegetation, dung, trampling or presence of animals). Field work was conducted from May to June 2019. The final field-dataset contained the above-mentioned environmental and species data for the 80 study plots comprising a total of 2308 records of 201 species.

### Reference data

To answer our study questions, we needed reference data from natural habitats of the same type as our study sites. In Europe, habitat types deemed important by the EU community through the Habitats Directive (Annex I, Council of the European Communities, 1992) – are considered more or less pristine nature with intact natural processes and hence generally the aim for natural habitat restoration projects. From the program for monitoring Danish Annex I habitats – the National Monitoring and Assessment Programme for the Aquatic and Terrestrial Environments (NOVANA) – we extracted plant species lists from 53,557 wetland plots covering all Denmark (Table 1). However, to put the restored plots into a broader context we also needed data from more degraded areas and plots with clear human-influence. Therefore, we also extracted plant species lists from 25,557 non-Annex-I wetland plots that were recorded through Danish municipalities’ inventories of natural and semi-natural protected nature. These plots encompassed: “improved meadows” (clear signs of earlier agricultural use, e.g. nutrient enrichment), “other meadows” (meadows with lower nutrient availability and more natural flora than improved meadows), “poor fens” (rather acidic nutrient poor fens, but not bogs or wet heathland), “nitrophile tall-herb communities” (e.g., reed swamps and riparian tall herb societies) and “nutrient poor meadows” (e.g., meadows affected by human activities but with no clear signs of artificial nutrient enrichment). Finally, we removed plots from both datasets having less than five species, lumped taxa to the species level and merged the two datasets resulting in a reference data set comprising 1,562,360 records of 1,425 vascular and bryophyte species covering 76,793 plots and 17 wetland habitat types (Table 1).

**Table 1.**
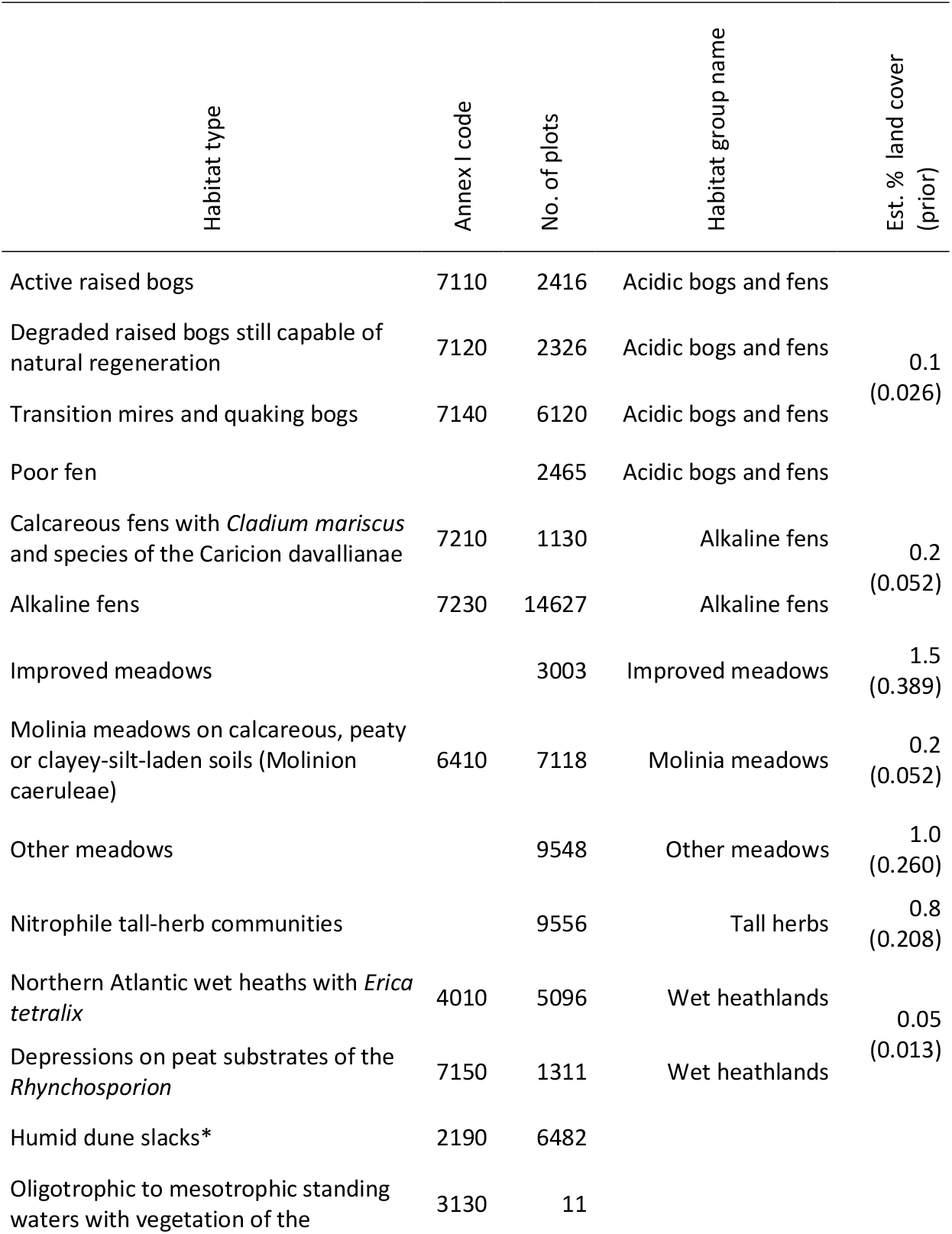
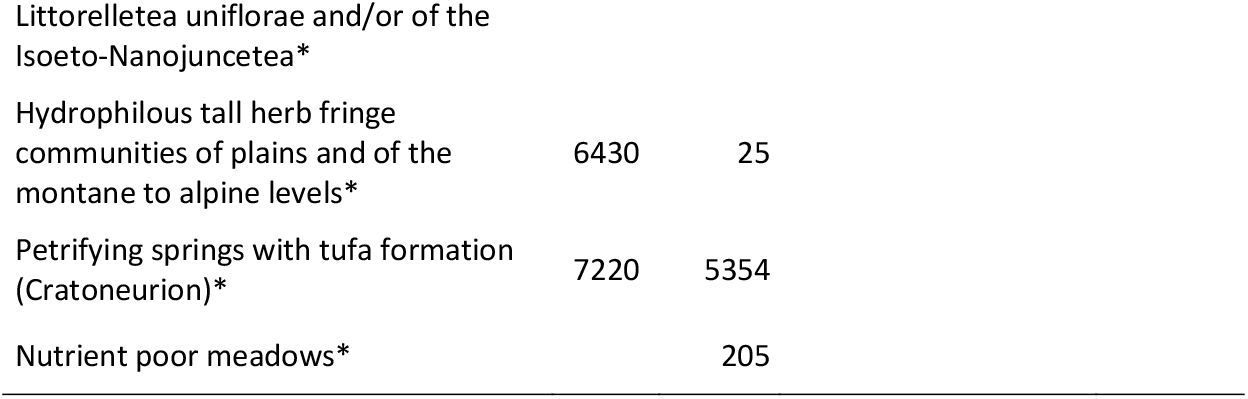
Overview of the habitat types included together with number of plots and the Annex I habitat code. Habitats within the same habitat group were lumped together for analysis. Habitats with poor coverage or lacking ecological meaning (i.e., not defined by the species living there) were left out of analysis (marked with *). The estimated land cover of each habitat group is given in the final column with priors used for quadratic discriminant analysis in parentheses (see main text).

### Data preparation

All vegetation plots used here, both those recorded as part of this study and the reference dataset were inventoried following the same method. The plots are 5-m radius circular plots inventoried in the growth season by botanical specialists. Prior to statistical analysis, the datasets were processed through several steps to add relevant information and to make sure data were consistent. Firstly, we renamed 22 species synonyms in the field-dataset to make names fully match those in the reference dataset. Then, we retrieved the updated Ellenberg Indicator Values (EIV) for plants from Hill et al. (1999) containing values indicating plants’ preferences for soil nitrogen (EIV_N_), reaction (EIV_R_), moisture (EIV_F_) and light availability (EIV_L_) covering 1,791 taxa. Species names were matched by use of the Taxonomic Name Resolution Service (Boyle et al., 2021, 100 names) combined with a manually check of synonyms for 52 species in the Danish species checklist (allearter.dk, accessed 4 October 2021). After lumping taxa in the EIV-dataset at the species level, we extracted indicator values for the 942 species in the EIV-dataset that were present in our field and reference datasets. For each plot in both datasets, we calculated the means of the four aforementioned indicator values and calculated the EIV_N_/EIV_R_-ratio based on the corresponding means. This ratio is a good indicator of actual soil fertility irrespective of the soil pH which tends to correlate with soil nitrogen (EIV_N_) (Andersen et al., 2013).

To enable using habitat-characteristic species in our statistical analysis, we retrieved a list of plant and byophyte species characteristic for alkaline fens from Andersen et al. (2013) (marked by “F” in their Appendix 1 table) and counted the number of such species in each plot in the field dataset. Since almost half of our study plots had zero alkaline fen indicator species, we also used a broader set of high-quality-habitat indicator species (Fredshavn & Ejrnæs, 2007, vascular plants, see species list in Table S1 in Supporting Information). These high-quality habitat indicator species are selected to indicate favorable conservation status cf. the Habitats Directive (Council of the European Communities, 1992) and include vascular plant species considered moderately to very sensitive to habitat destruction (Fredshavn & Ejrnæs, 2007).

### Statistical analysis

#### Habitat classification

To enable comparison between our field and reference data, we created a classification model for classifying plots into habitat types based on environmental conditions (Ellenberg Indicator Values, see above). This classification model was based on the reference dataset. However, to construct a biologically sound classification model, we first excluded plots with poor habitat representation (Table 1). We also excluded plots belonging to habitat types with poor ecological meaning (i.e., not defined by the species living there): Petrifying springs (EU habitat code: 7220) are defined by the hydrology and dune slacks (habitat code: 2190) are defined by topographic position and can vary considerably in soil moisture, soil nutrient avilability and successional stage (Table 1). Secondly, we grouped plots belonging to habitats with more or less the same species communities (see Table 1). Then, we ran a quadratic discriminant analysis (QDA) to predict these grouped habitat types from the plants’ overall preferences for nitrogen (plot mean EIV_N_), soil reaction (EIV_R_), moisture level (EIV_F_) and light availability (EIV_L_, Fig. 2). Since the number of plots in the reference data set for each habitat group does not reflect their actual abundance in the landscape, we used priors weighted accordingly, to avoid overfitting to habitat groups that are overrepresented in the data set, e.g., alkaline fens. Table 1 contains the estimated national land cover percentage of each habitat group (Annex I habitat coverage from Fredshavn et al., 2019, and the rest from supplementary table 1 in Ejrnæs et al., 2018) and the priors used in QDA.

**Figure 2.**
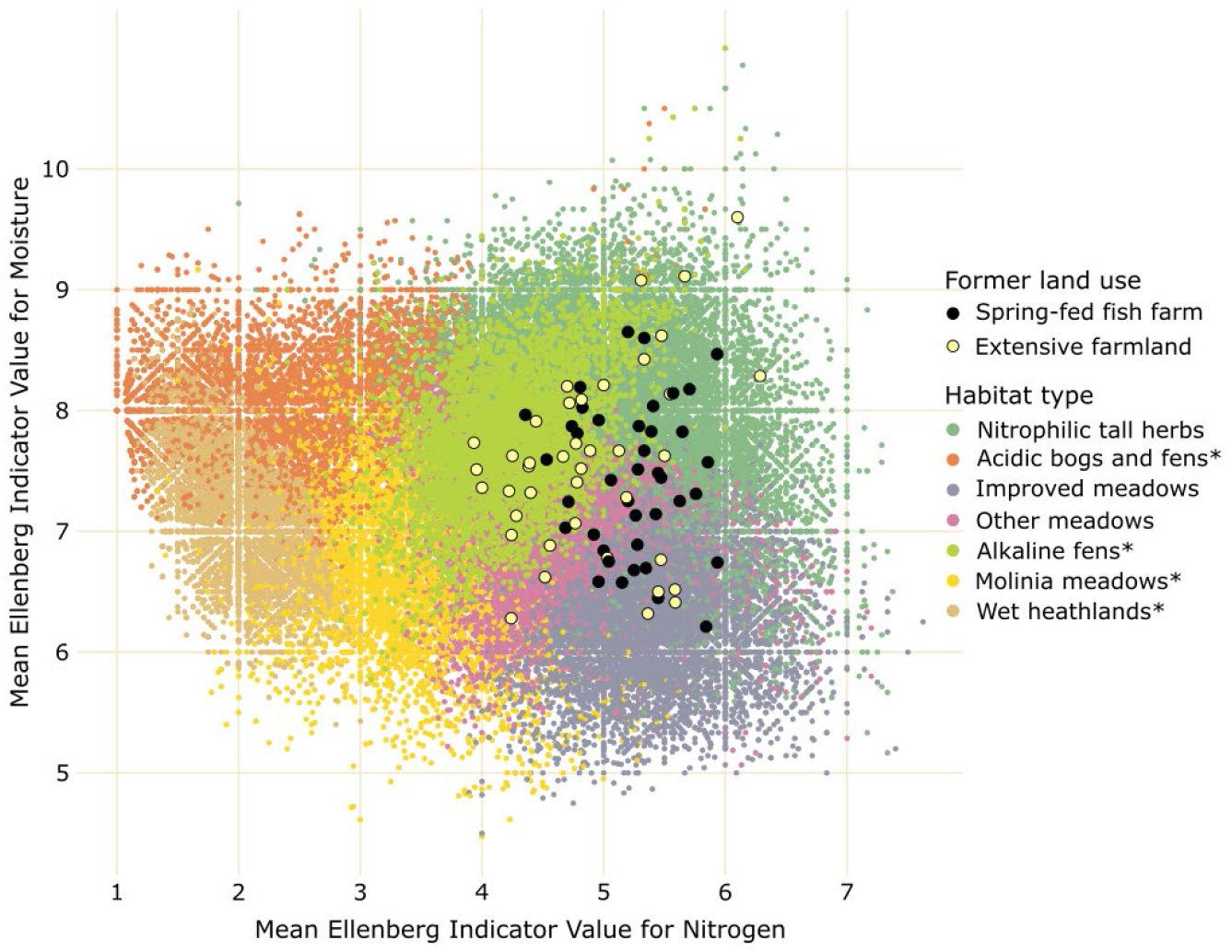
The predicted distribution of reference plots (small points) and study plots (larger points) along the moisture and nutrient gradients in this study (see main text). The different wetland types in the legend are explained in the main text. Habitat types marked with asterisks are Annex I habitats in the European Habitats Directive (Council of the European Communities, 1992).

Subsequently, we used the classification model to predict which habitat types were most likely for the restored wetland plots in our study. In all cases but two, our study plots had highest probability (posterior) for being alkaline fens (Fig. 3). Hence, we assumed in the remaining analyses that the restored wetlands are moving towards becoming alkaline fens, so we considered this the main target habitat type.

**Figure 3.**
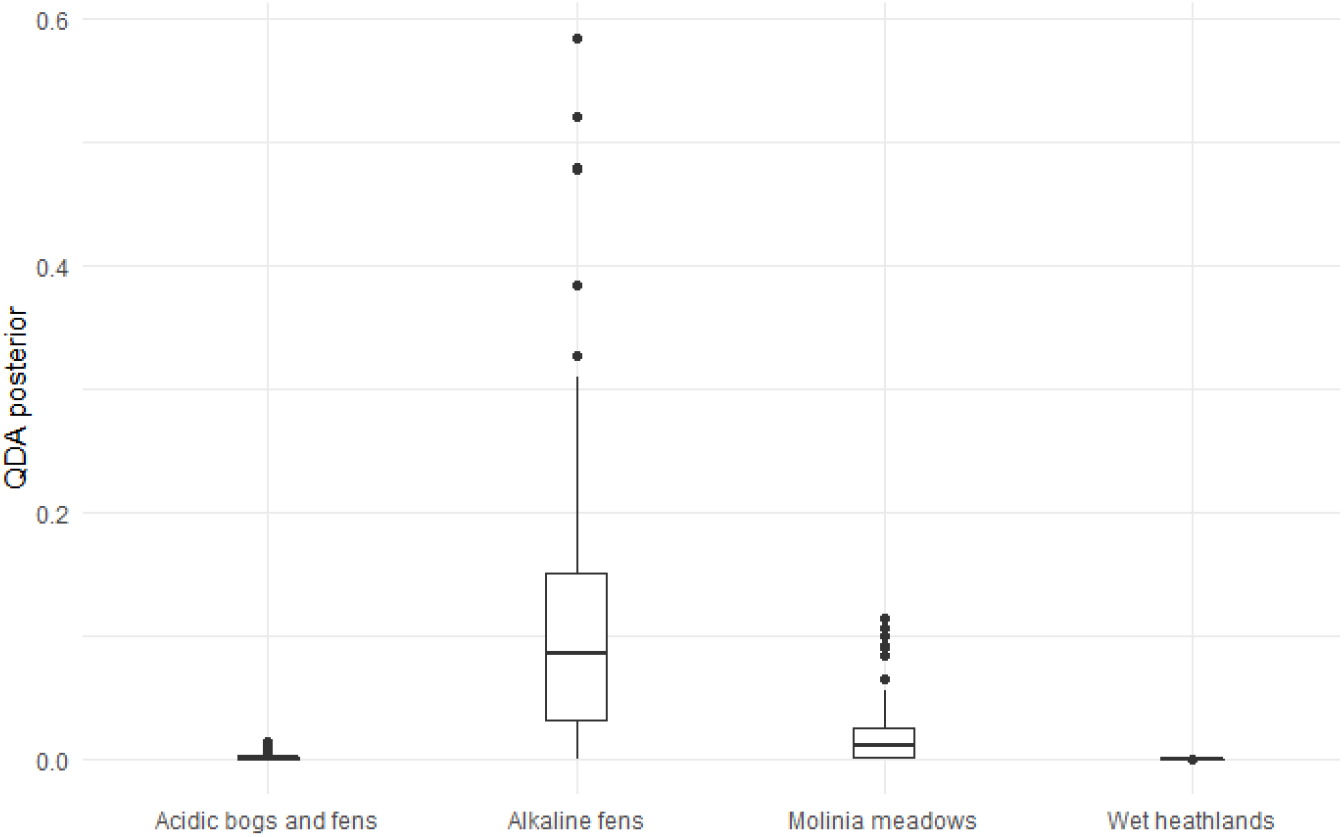
Posteriors from the Quadratic Discriminant Analysis (QDA) described in the main text for the habitats used in this study and listed in Annex I in the European Habitats Directive (Council of the European Communities, 1992).

#### Variables for models and analyses

##### Response variables

To investigate the effects of restoration practice – such as rewetting, grazing and time – on restoration success, we constructed four statistical models with four different response variables. Firstly, to analyse these factors in relation to how close a given restoration plot is to Annex I wetlands we used the posteriors from the QDA as response variables; (1) both the posteriors for alkaline fens only (i.e., the probability that a restored plot is an Annex I alkaline fen, coded 7230) and (2) the summed posteriors (i.e., the probability that a restored plot is any of the included Annex I wetland habitats). Secondly, we used (3) the number of high-quality-habitat indicator species as response instead. These species are considered vulnerable to habitat deterioration by agricultural intensification (Fredshavn & Ejrnæs, 2007) such as tilling, fertilization, drainage and cropping (e.g. grass leys). Thirdly, we used (4) the mean EIV_N_/EIV_R_-ratio as response variable in a fourth model. We expected that analysing this variable could give insight into how soil nutrient availability varies under different restoration settings.

##### Explanatory variables

To investigate the effect of *former land-use*, we included a binary explanatory variable denoting whether a study site (and thereby the four plots within it) was either a restored former fish farm in a spring dominated area or a former extensive farmland where drainage is now reduced as part of the wetland restoration (see further explanation in the section “Study area”). To test the effect of time, we used the variable *years since restoration* giving the years elapsed since the completion of the restoration projects. To study effects of grazing by large herbivores, we created a variable representing *grazing* conditions: as we did not have information of grazing pressure, we used the grazing information as a binary variable; plots with signs of grazing (see above) were marked as grazed and others ungrazed. To analyse the effect of rewetting, i.e., to which degree the restored hydrology corresponds to historic hydrological conditions, we included rewetting in our analyses. The effect of rewetting is known to be spatial-scale dependent (Rolls et al., 2018) and therefore we calculated two rewetting variables to make sure we covered the likely spatial scales at which hydrology can affect local plant diversity; one reflecting conditions at *plot-scale* and one at *study-site-scale*. To do this we digitalized historical hydrological conditions at our study sites by delineating areas that used to be wet, moist, and dry in the first large-scale mapping of Danish territory in 1862– 1899 (“Høje målebordsblade”, freely available through www.dataforsyningen.dk). Recall that in the field, we mapped the corresponding current wetness of each site. For all study plots, we subsequently determined the wetness of the majority of the plot (recall: plots are 5-m radius circles) historically and currently. For example, if 60 % of the plot was wet and 40 % was moist, we considered the plot wet in our statistical analyses. We constructed a binary *plot-scale* rewetting variable holding a yes if a plot was as wet or wetter than historically, otherwise a no. At *site-scale*, we calculated the areal percentage that were as wet or wetter than it used to be historically. For this calculation, we did not consider areas which were dry historically as this study concerns restoration of wetlands. We weighted these percentages for each wetness-category by its original area and then summed them. The result was a variable reflecting to what degree historically moist or wet study sites are contemporarily at least as moist or wet as they were in the late 1800s. In our modelling (see below), the site-scale rewetting consistently gave better models (Akaike’s Information Criterion, AIC, Akaike, 1974) and the plot- and site-scale variables gave the same results, so we did not use the plot-scale rewetting variable in our final models.

#### Comparing restored and reference wetlands

To gain insight into the floristic differences between restored wetland sites and natural alkaline fens (recall that the restored sites had highest probability of being alkaline fens), we compared the number of characteristic alkaline fen species in our study plots and in the alkaline fen reference plots using a Wilcoxon rank sum test.

To investigate the degree to which soil fertility influences restoration success, we tested if our study plots held as many alkaline fen characteristic species as we would expect from the prevailing soil fertility. To do this we constructed a Generalized Linear Model (GLM, poisson distributed errors) with number of characteristic alkaline fen species (see above) as response and mean EIVN/EIV_R_-ratio as explanatory variable. This model included only alkaline fen plots from our reference dataset (recall that these are the most natural alkaline fens in the country, see above). Then we used it to predict the number of characteristic alkaline fen species in our restoration plots and finally that number was compared to the actual observed number of these species by a Wilcoxon rank sum test. To avoid circularity in this analysis, we used a mean EIV_N_/EIV_R_-ratio calculated without the characteristic alkaline fen species.

#### Modelling the effects of nutrients, rewetting, grazing and time

For each of the four response variables, we constructed a GLM having errors following either a binomial (QDA posteriors), poisson (no. of indicator species) or gaussian distribution (EIVN/EIVR-ratio) to test the effects of (1) former land-use, (2) number of years since restoration, (3) presence or absence of grazing, (4) the degree to which hydrology was restored to historical conditions and (5) the mean EIVN/EIVR-ratio. The latter was only used in the model of indicator species, as the QDA was based on EIVs, and hence using it in the other models would cause circularity. In the modelling of posteriors from the QDA, we used a two-column matrix format having no. of successes and no. of failures as columns. To create this response-variable format, we used the posteriors in integer percent as “no. of successes” and 100 % - “no. of successes” as “no. of failures”.

We used a stepwise backwards model selection procedure based on AIC (Akaike, 1974), removing the variable causing the highest drop in AIC in each step and stopping when ΔAIC did not change more than 2 (Burnham & Anderson, 2002). As part of the model selection procedure, we tested all two-way interactions and kept those that improved the model significantly (same ΔAIC criteria as above). As previous studies have shown that the effect of time after restoration is not necessarily linear (Matthews et al., 2009b), we also included a quadratic term of this variable to allow for a non-linear fit. After modelling we checked all models’ residuals for spatial autocorrelation using Moran’s I (Paradis, 2015), and found none (P >> 0.05). We checked the final models for model misfit by inspecting predicted vs. actual residual plots, and we checked that the poisson model was not overdispersed.

## Results

Our results showed that the probability that a restored wetland is an Annex I wetland habitat (including alkaline fens) is generally below 10 % and only four plots reached a probability of more than 50 % (Fig. 3). Also, we only found half as many characteristic alkaline fen species in restored wetlands (median = 1 species) than in reference fens (median = 2 species, Wilcoxon rank sum test P-value ≪ 0.001). In addition to this, we demonstrated that the number of characteristic alkaline fen species did not deviate from what could be expected under the prevailing soil nutrient availability in the restored sites (Wilcoxon rank sum test P-value = 0.17).

Our two models of probabilities (i.e., QDA posteriors) that a given plot is either an Annex I alkaline fen or any Annex I wetland habitat were almost identical and explained about 18–19 % of the variation in data. Therefore, we report and discuss the results of the alkaline fen model only, since almost all plots were closest to this habitat type. Results for the model for all Annex I wetlands can be found in Fig S1.

Generally, the plots’ probability of being alkaline fen wetlands were higher in former extensive farmland compared to former spring-fed fish farms (Table 2) notably in non-grazed plots (Fig. 4A). The effect of time since restoration was the most important factor for this probability (Table 2) and peaked after c. 12–15 years in grazed areas and after c. 30 years in non-grazed areas (Fig 4B) and was highest in the plots with poor rewetting (Fig 4D). Generally, rewetting had a negative influence on the probability of a plot being a alkaline fen wetland habitat, but this was most pronounced in former extensive farmlands (Fig. 4C). Full modelling results are shown in Table 3. All significant interactions are shown in Fig. 4.

**Figure 4.**
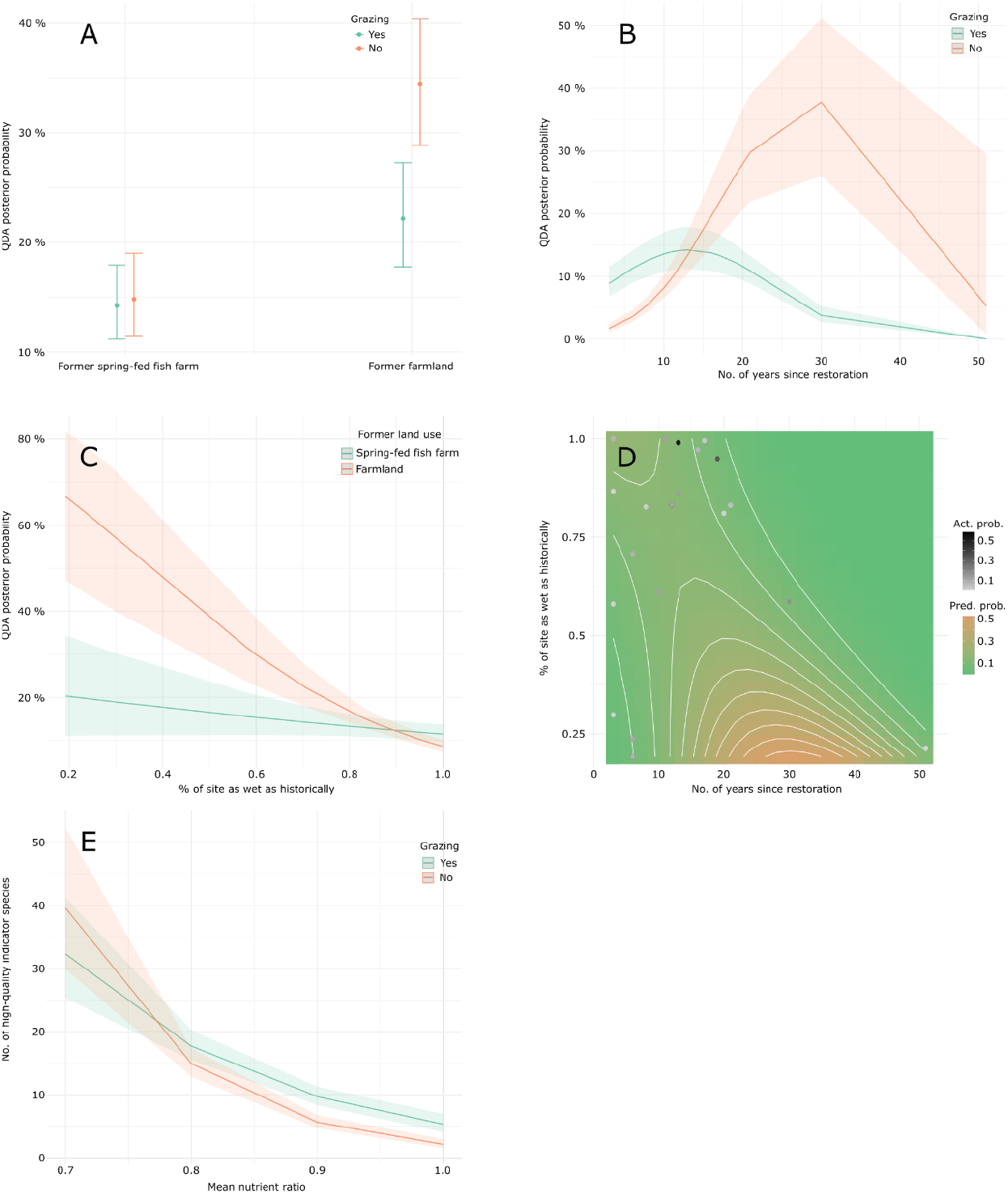
Predicted effects of the interactions and squared terms in the models of (1) the probability of being an alkaline fen (A, B, C, D) and (2) the number of high-quality habitat indicator species (E). Where uncertainty is shown this corresponds to the 95% confidence interval. Panel D shows a contour plot of the interaction between two continuous variable and hence the contours correspond to the predicted Quadratic Discriminant Analysis (QDA) posterior probability (Pred. prob.) shown in the Y axes of the other graphs. A contour plot extrapolates the data and hence is invalid far outside the areas where the actual probabilities (Act. prob.) for each plot is shown by dots (e.g., the upper right corner of the plot).

**Table 2.** Explanatory variable importances given as the residual deviance after the explanatory variable in question has been removed from the model. The higher the residual deviance after variable removal the more important is the explanatory variable considered. In cases with interactions, these also count in the importance (i.e., interactions with the variable in question was also removed from model for calculation of variable importance). For comparison, model null deviance and residual deviance is given along with deviance explained and the number of observations used in the model (N).

| Explanatory variable | Prob. of<br>alkaline fen | Prob. of Annex I<br>wetland | # high-quality<br>habitat<br>species |
| --- | --- | --- | --- |
| Former land-use | 923.0 | 1131.6 | 105.0 |
| Grazing | 917.8 | 1092.6 | 115.2 |
| # years since rest. | 932.3 | 1111.4 | 104.8 |
| # years since rest.<sup>2</sup> | 853.4 | 1030.5 |  |
| Area % wetter than before | 881.9 | 1064.3 |  |
| Former land-use * Grazing | 835.5 | 1014.3 |  |
| Grazing * # years since rest. | 915.0 | 1090.0 |  |
| Former land-use * Total area % wetter<br>than before | 861.9 | 1045.1 |  |
| # years since rest. * Area % wetter than<br>before | 863.6 | 1042.2 |  |
| Mean N/R ratio |  |  | 256.8 |
| Grazing * Mean N/R ratio |  |  | 104.5 |
| Null deviance | 1016.5 | 1225.4 | 274.0 |
| Full model deviance | 825.0 | 1004.1 | 94.1 |
| Deviance explained | 18.8 % | 18.0 % | 65.7 % |
| N | 76 | 76 | 76 |

Secondly, our results showed that soil nutrient availability was the most important factor for the number of high-quality habitat indicator species (Table 2) with a general negative impact on this richness. The effect of grazing was most pronounced in the most nutrient rich plots and here it had a positive effect on the number of high-quality habitat indicator species (Fig 4E) while time since restoration generally had a negative influence (Table 3). Figure 5 shows two examples of study plots with a high number and two with a low number of high-quality habitat indicator species and their soil nutrient availability and grazing status.

**Figure 5.**
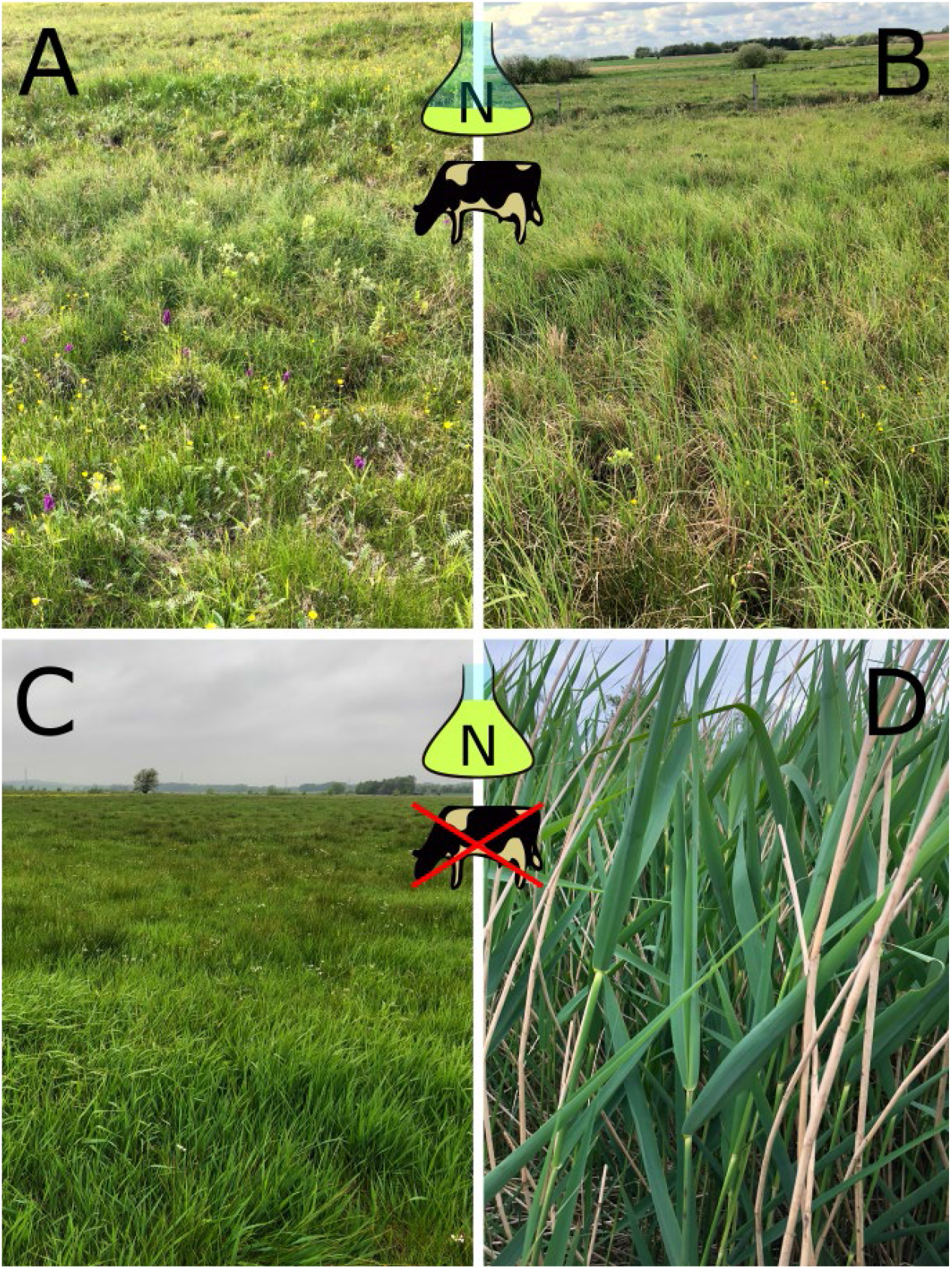
Two plots with a high number of high-quality habitat indicator species (14 (A) and 27 (B) species) and two with a low number of these species (2 (C) and 1 (D) species). Full and almost empty lab-bottles indicate high and low soil nutrient availability (N) while ungrazed or grazed plots are indicated with a cow with or without a red cross respectively.

**Table 3.** Modelling results overview. The three main groups of columns represent results for each of the three models having the given response variable.

| Explanatory variable | Prob. of alkaline fen |  |  | Prob. of Annex I wetland |  |  | # high-quality habitat species |  |  |
| --- | --- | --- | --- | --- | --- | --- | --- | --- | --- |
|  | log(OR) | 95% CI | p | log(OR) | 95% CI | p | log(IRR) | 95% CI | p |
| Former land-use |  |  |  |  |  |  |  |  |  |
| *Spring-fed fish farm* | — | — |  | — | — |  | — | — |  |
| *Extensive farmland* | 2.6 | 1.8, 3.5 | **<0.001** | 2.7 | 1.9, 3.5 | **<0.001** | -0.27 | -0.43, -0.11 | **<0.001** |
| Grazing |  |  |  |  |  |  |  |  |  |
| Yes | — | — |  | — | — |  | — | — |  |
| No | -2.2 | -2.8, -1.7 | **<0.001** | -2.0 | -2.5, -1.5 | **<0.001** | 2.8 | 0.95, 4.7 | **0.003** |
| # years since rest. | 0.38 | 0.25, 0.51 | **<0.001** | 0.34 | 0.22, 0.46 | **<0.001** | -0.01 | -0.02, 0.00 | **0.001** |
| # years since rest.<sup>2</sup> | -0.01 | -0.01, 0.00 | **<0.001** | 0.00 | -0.01, 0.00 | **<0.001** |  |  |  |
| % of site as wet as historically | 3.9 | 2.6, 5.2 | **<0.001** | 3.7 | 2.5, 4.9 | **<0.001** |  |  |  |
| Former land-use * Grazing |  |  |  |  |  |  |  |  |  |
| *Extensive farmland \* No* | 0.56 | 0.22, 0.91 | **0.001** | 0.50 | 0.19, 0.82 | **0.002** |  |  |  |
| Grazing * # years since rest. |  |  |  |  |  |  |  |  |  |
| *No \* # years since rest.* | 0.17 | 0.13, 0.20 | **<0.001** | 0.15 | 0.11, 0.18 | **<0.001** |  |  |  |
| Former land-use * % of site as wet as historically |  |  |  |  |  |  |  |  |  |
| *Extensive farmland \* % of site as wet as historically* | -2.9 | -3.9, -2.0 | **<0.001** | -2.9 | -3.8, -2.0 | **<0.001** |  |  |  |
| # years since rest. * Area % wetter than before | -0.35 | -0.46, -0.23 | **<0.001** | -0.31 | -0.42, -0.21 | **<0.001** |  |  |  |
| Mean N/R ratio |  |  |  |  |  |  | -6.0 | -7.5, -4.5 | **<0.001** |
| Grazing * Mean N/R ratio |  |  |  |  |  |  |  |  |  |
| *No \* Mean N/R ratio* |  |  |  |  |  |  | -3.7 | -6.0, -1.5 | **0.001** |

OR = Odds Ratio, CI = Confidence Interval, IRR = Incidence Rate Ratio, Prob. = Probability, # = number of, red. = reduced, drain. = drainage, Mean N/R ratio = the ratio between mean Ellenberg Indicator Values for soil nitrogen(N) and soil pH (R), a measure of soil nutrient availability.

Thirdly, we found that the mean soil nutrient availability in the restored wetlands was only affected by the former land-use with former extensive farmlands having a lower nutrient availability (β = - 0.04, P = 0.010) than former spring-fed fish farms.

## Discussion

In this study, we showed that restored wetlands generally hold a vegetation that is far from reference Annex I wetlands – both with regard to characteristic species and the more general plant species composition – and this was also true for sites where restoration took place > 20 years ago. Our results confirm several previous findings reporting the same tendency (Mälson et al., 2008; Matthews et al., 2009b; Baumane et al., 2021). We also showed that relatively nutrient rich plots had significantly lower richness of high-quality habitat indicator species and that – given the prevailing nutrient availability – the current richness of characteristic alkaline fen species corresponds to predictions based on a large reference dataset. This suggests that the main barrier to restoration success is nutrient load rather than dispersal or establishment limitation. If restoration of naturally infertile soils is not ensured, we predict that Annex I wetland habitats in stream valleys cannot be restored successfully and instead the areas will likely grow into species poor wetlands with dominance of highly competitive plant species (Naja et al., 2017). An effective way to achieve a natural nutrient balance is to remove the nutrient-rich topsoil possibly combined with addition of lime where appropriate (Mälson & Rydin, 2007; Mälson et al., 2008). In fact, on former farmland complete topsoil removal is sometimes the only way to achieve natural nutrient conditions in wetland restoration (Smith et al., 2011).

Time since restoration had an overall positive influence on the probability of a study plot being an alkaline fen, but only in the first 12–30 years after restoration. In grazed areas this effect was significantly lower, and the effect of time wore off earlier. One possible explanation for this pattern is that some plant species characteristic in alkaline fens – e.g., *Carex nigra* (L.) Reichard, *Lychnis flos-cuculi* L. and *Menyanthes trifoliata* L. – may establish rather quickly making the vegetation resemble a poorly developed alkaline fen. However, after some years – if the nutrient load is too high as indicated in this study (see above) – the characteristic alkaline fen species could disappear again due to competitive exclusion (Borer et al., 2014). Matthews et al. (2009b) found a similar pattern for several wetland-restoration success-indices including one based on plant species’ conservation value. They also considered invasion of tall growing herbs a few years after restoration a plausible explanation for this finding. Additionally, this could explain why we found a small but significant negative effect of time on the number of high-quality habitat indicator species. Such a phenomenon could be aggravated by finishing wetland restoration with a nutrient rich topsoil layer and seeding of agricultural grass cultivars, a common practice in older wetland restoration projects in Denmark but now less frequent (e.g., Andersen et al., 2005). These cultivars are highly competitive and quickly establishes a dense monospecific vegetation. Another explanation could be that the oldest restoration projects may have been undertaken differently than the younger ones, with less focus on creating optimal conditions for wetland plant diversity. In a recent study, Baumane et al. (2021) found no effect of time since restoration on the vegetation in riparian wetlands. They suggested this to be a consequence of too high nutrient input and possibly dispersal limitation. We do not find evidence that dispersal limitation is causing the observed development of restored wetlands because the number of characteristic alkaline fen species do not deviate from the expected number given environmental conditions. Our results suggest that we cannot expect any further development into more characteristic wetland nature with time (see discussion above on nutrients).

Our study showed that previous land-use matters for restoration success. Contrary to our expectations, the probability of a plot being an alkaline fen was higher in former extensive farmlands than in former spring-fed fish farms. However, when analyzing nutrient differences between the two land-use types we learned that our former fish-farms had higher soil nutrient availability than the former extensive farmlands, possibly explaining why these former extensive agricultural areas appeared more like alkaline fens than the former fish farms. In addition to this, when we controlled for soil nutrient availability in our analysis of high-quality indicator species this pattern changed; the number of these species were higher in former spring-fed fish farms than in former farmland areas. This points to that – soil nutrient availability being at equal levels – restoration of alkaline fens is probably most effective in sites where clean groundwater exfiltrates diffusely from the underground and possibly washes away nutrients and in some cases immobilizes the phosphor pool (Boomer & Bedford, 2008; Venterink et al., 2009; Audet et al., 2015). In our study sites, results indicate that the nutrient rich sludge from the fish-farms was not consistently removed or that the fish-farms received nutrient rich soil during restoration. Finally, there may be other differences in restoration practice between the two former land-uses, for example whether practitioners attempted to restore natural microtopography which is known to be important for natural wetland plant diversity (Moeslund et al., 2013).

Grazing by cattle or horses had a positive effect on the probability of a plot being an alkaline fen or an Annex I wetland habitat, but only in the first years after restoration. We also found a positive effect of grazing on the number of high-quality habitat indicator species, but only in the most nutrient-rich parts of our study sites. On the other hand, grazing did not improve the probability of a plot being an alkaline fen in the later time periods after restoration, but this negative effect was only clear in former extensive farmland areas and was relatively small in the oldest sites. Generally grazing is considered important for freshwater wetland biodiversity as it creates heterogenous vegetation and terrain and thereby numerous microhabitats for specialist species (Biró et al., 2019, and references therein), and our study also partly supports this. The lack of a consistent clear positive effect of grazing may also reflect the low precision of our grazing data, possible confounding with other activities such as sward improvements or perhaps an overriding effect of eutrophication.

The degree to which an area was rewetted to resemble historical wetness generally had a negative impact on the probability of a plot being an alkaline fen, notably in former extensive farmlands. This is counterintuitive because clearly, restoring hydrology must have a positive impact on wetland restoration as these habitat types cannot be restored without water (Pfadenhauer & Grootjans, 1999; Mälson et al., 2008). In wetlands, different species have specialized in different hydrological niches, and these niches can only be fully present if the hydrology is intact (Rolls et al., 2018). However, here we only see such a positive effect of rewetting in the first c. 10–15 years. Based on our results (see discussion about nutrients above), we suggest that this could be a consequence of too high a soil or water nutrient availability in the restored sites generally. In other words, we believe the most plausible explanation is that the sites benefit from successful rewetting in the first few years and then the nutrient balance short-circuits this positive development and becomes the main determinant of the final restoration outcome.

### Mertainties

In our models, the results regarding rewetting and time since restoration comes with uncertainties. Determining contemporary wetness in the field and subsequently historical wetness based on old maps is a difficult task and the resulting variables will probably vary slightly depending on the person undertaking this task. Here, we carefully conducted this task and only one person was involved in the historic map interpretations to ensure consistency. Also, we believe that site-scale hydrology acts at a relatively large spatial scale further reducing the impact by the uncertainties associated with this variable. Regarding time since restoration, we only had three sites (12 plots) that were older than 20 years and this clearly causes some uncertainty regarding the effect of time in the later years after restoration. That said, we believe our results are rather solid and trustworthy.

## Acknowledgements

We sincerely thank Sebastian Niebuhr McQueen and Nanna Emilie Hesthaven Mikkelsen for help with fieldwork. We also thank the people involved in LIFE IP Natureman who helped us with data and knowledge about restoration projects. This study is a contribution to the project “Mapping, restoration and management of groundwater-fed fens and springs” funded by Aage V. Jensen Naturfond.

