## Supporting Information for "High nutrient loads hinder successful restoration of natural habitats in freshwater wetlands"

**Table S1**. List of the high-quality habitat indicator species from (Fredshavn and Ejrnæs 2007) used in the study.

| |
|---|
| Ajuga reptans |
| Alisma plantago-aquatica |
| Anemone nemorosa |
| Angelica sylvestris |
| Anthoxanthum odoratum |
| Avenula pubescens |
| Briza media |
| Caltha palustris |
| Cardamine amara |
| Cardamine pratensis |
| Carex canescens |
| Carex cespitosa |
| Carex diandra |
| Carex echinata |
| Carex flacca |
| Carex leporina |
| Carex nigra |
| Carex oederi |
| Carex panicea |
| Carex paniculata |
| Carex rostrata |
| Centaurea jacea |
| Chrysosplenium oppositifolium |
| Cicuta virosa |
| Cirsium palustre |
| Comarum palustre |
| Cynosurus cristatus |
| Dactylorhiza incarnata |
| Dactylorhiza majalis |
| Dryopteris carthusiana |
| Dryopteris dilatata |
| Eleocharis palustris |
| Eleocharis uniglumis |
| Epilobium palustre |
| Epilobium parviflorum |
| Equisetum fluviatile |
| Equisetum palustre |
| Eriophorum angustifolium |
| Galium boreale |
| Galium palustre |
| Galium uliginosum |
| Geum rivale |
| Hydrocotyle vulgaris |
| Hypericum tetrapterum |

| |
|---|
| Iris pseudacorus |
| Juncus articulatus |
| Lotus pedunculatus |
| Luzula campestris |
| Luzula multiflora |
| Lychnis flos-cuculi |
| Lysimachia thyrsiflora |
| Menyanthes trifoliata |
| Myosotis laxa |
| Myosotis scorpioides |
| Pedicularis palustris |
| Peucedanum palustre |
| Potentilla erecta |
| Primula veris |
| Prunella vulgaris |
| Ranunculus flammula |
| Ranunculus lingua |
| Rhinanthus minor |
| Rumex hydrolapathum |
| Sagina nodosa |
| Salix hastata |
| Salix pentandra |
| Salix repens |
| Schoenoplectus tabernaemontani |
| Scirpus sylvaticus |
| Stachys palustris |
| Stellaria alsine |
| Stellaria graminea |
| Stellaria palustris |
| Succisa pratensis |
| Triglochin palustris |
| Valeriana sambucifolia |
| Veronica anagallis-aquatica |
| Veronica beccabunga |
| Veronica montana |
| Vicia cracca |
| Viola palustris |

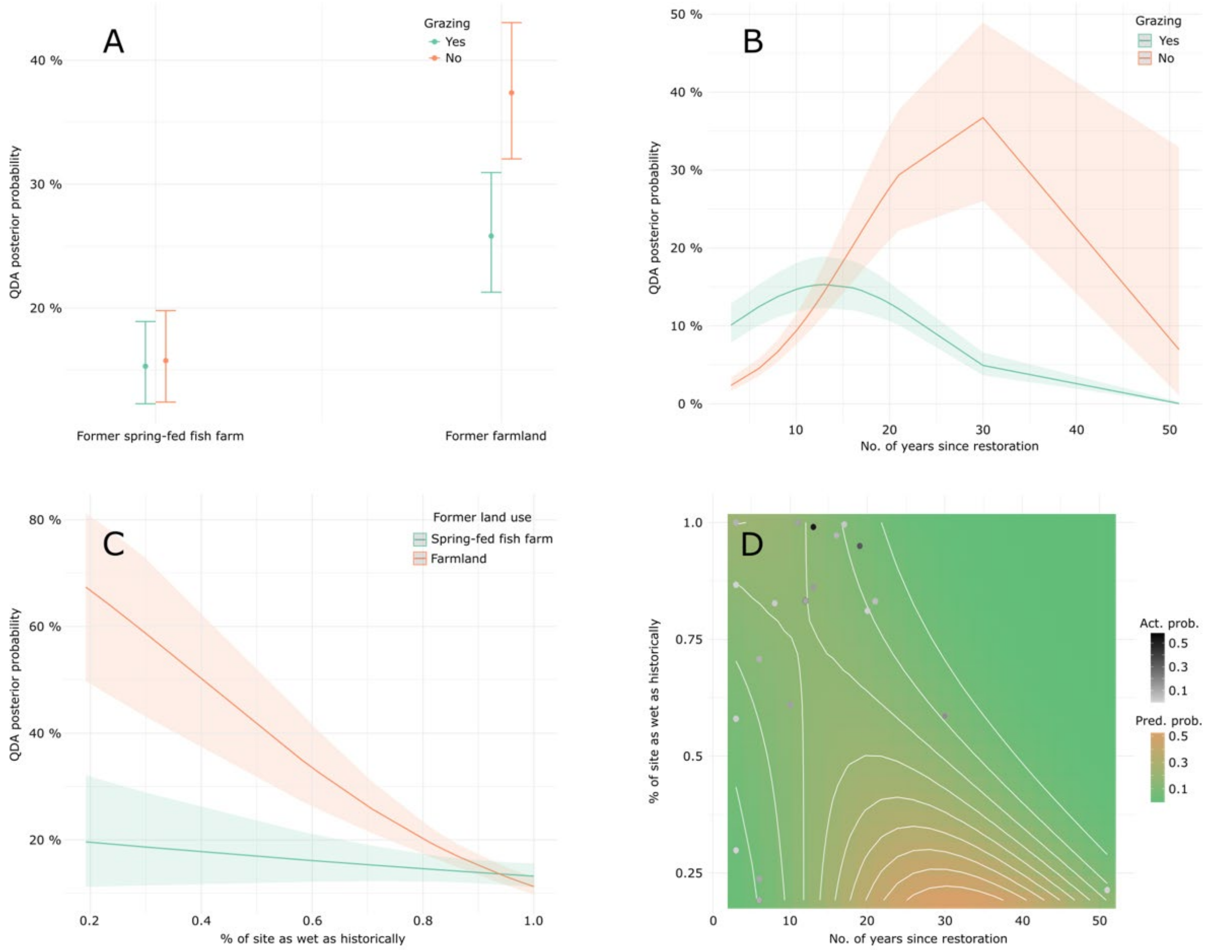


**Figure S1**. Predicted effects of the interactions and squared terms in the models of (1) the probability of being any Annex I wetland habitat type (A, B, C, D). Where uncertainty is shown this corresponds to the 95% confidence interval. Panel D shows a contour plot of the interaction between two continuous variable and hence the contours correspond to the predicted QDA posterior probability (Pred. prob.) shown in the Y axes of the other graphs. A contour plot extrapolates the data and hence is invalid far outside the areas where the actual probabilities (Act. prob.) for each plot is shown by dots (e.g., the upper right corner of the plot).
